# One enzyme reverse transcription qPCR using Taq DNA polymerase

**DOI:** 10.1101/2020.05.27.120238

**Authors:** Sanchita Bhadra, Andre C. Maranhao, Andrew D. Ellington

## Abstract

Taq DNA polymerase, one of the first thermostable DNA polymerases to be discovered, has been typecast as a DNA-dependent DNA polymerase commonly employed for PCR. However, Taq polymerase belongs to the same DNA polymerase superfamily as the Molony murine leukemia virus reverse transcriptase and has in the past been shown to possess reverse transcriptase activity. We report optimized buffer and salt compositions that promote the reverse transcriptase activity of Taq DNA polymerase, and thereby allow it to be used as the sole enzyme in TaqMan RT-qPCR reactions. We demonstrate the utility of Taq-alone RT-qPCR reactions by executing CDC SARS-CoV-2 N1, N2, and N3 TaqMan RT-qPCR assays that could detect as few as 2 copies/µL of input viral genomic RNA.

## INTRODUCTION

RT-qPCR remains the gold standard for the detection of SARS-CoV-2. However, the complexity of this assay sometimes limits its use in either resource-poor settings or in circumstances where reagent availability has become limited. To overcome these limitations, we have previously advocated the use of a thermostable reverse transcriptase (RT) / DNA polymerase (DNAP), which we have termed RTX (and which is distinct from RTx, from New England Biolabs) for use as the RT component of RT-qPCR. RTX is an evolved variant of the high-fidelity, thermostable DNA polymerase from *Thermococcus kodakaraensis* (KOD DNAP) with relaxed substrate specificity allowing it to perform as both a DNA- and RNA-directed DNA polymerase^*1*^. RTX has been shown to serve as the RT component in standard TaqMan probe-based RT-qPCR and as the sole enzyme component for dye-based RT-qPCR (https://www.biorxiv.org/content/10.1101/2020.03.29.013342v4).

The basis for the directed evolution of RTX is that many DNA polymerases have reverse transcriptase activity, with some of them, such as the polymerase from *Thermus thermophilus* (Tth), having fairly substantial activity^*2*^. This is a discovery that appears to resurface every so often, but that can be traced back to 1989^*3*^. The use of a single enzyme for both reverse transcription and DNA polymerization can simplify molecular diagnostic assays, including isothermal amplification assays^*4*^. In the midst of reagent supply issues, this led us to wonder whether it might prove possible to use readily available thermostable DNA polymerases as single enzyme solutions for RT-qPCR. We find that a relatively simple buffer (“Gen 6 A”) allows robust detection of SARS-CoV-2 in TaqMan probe-based RT-qPCR. We have validated this buffer mix with Taq DNAP from several commercial sources.

## METHODS

### Chemicals and reagents

All chemicals were of analytical grade and were purchased from Sigma-Aldrich (St. Louis, MO, USA) unless otherwise indicated. All enzymes and related buffers were purchased from New England Biolabs (NEB, Ipswich, MA, USA), Thermo Fisher Scientific (Waltham, MA, USA), or Promega (Madison, WI, USA) unless otherwise indicated. All oligonucleotides and TaqMan probes (**Table 1**) were obtained from Integrated DNA Technologies (IDT, Coralville, IA, USA). SARS-CoV-2 N gene armored RNA was obtained from Asuragen (Austin, TX, USA). SARS-CoV-2 viral genomic RNA was obtained from American Type Culture Collection (Manassas, VA, USA).

**Table 1.**
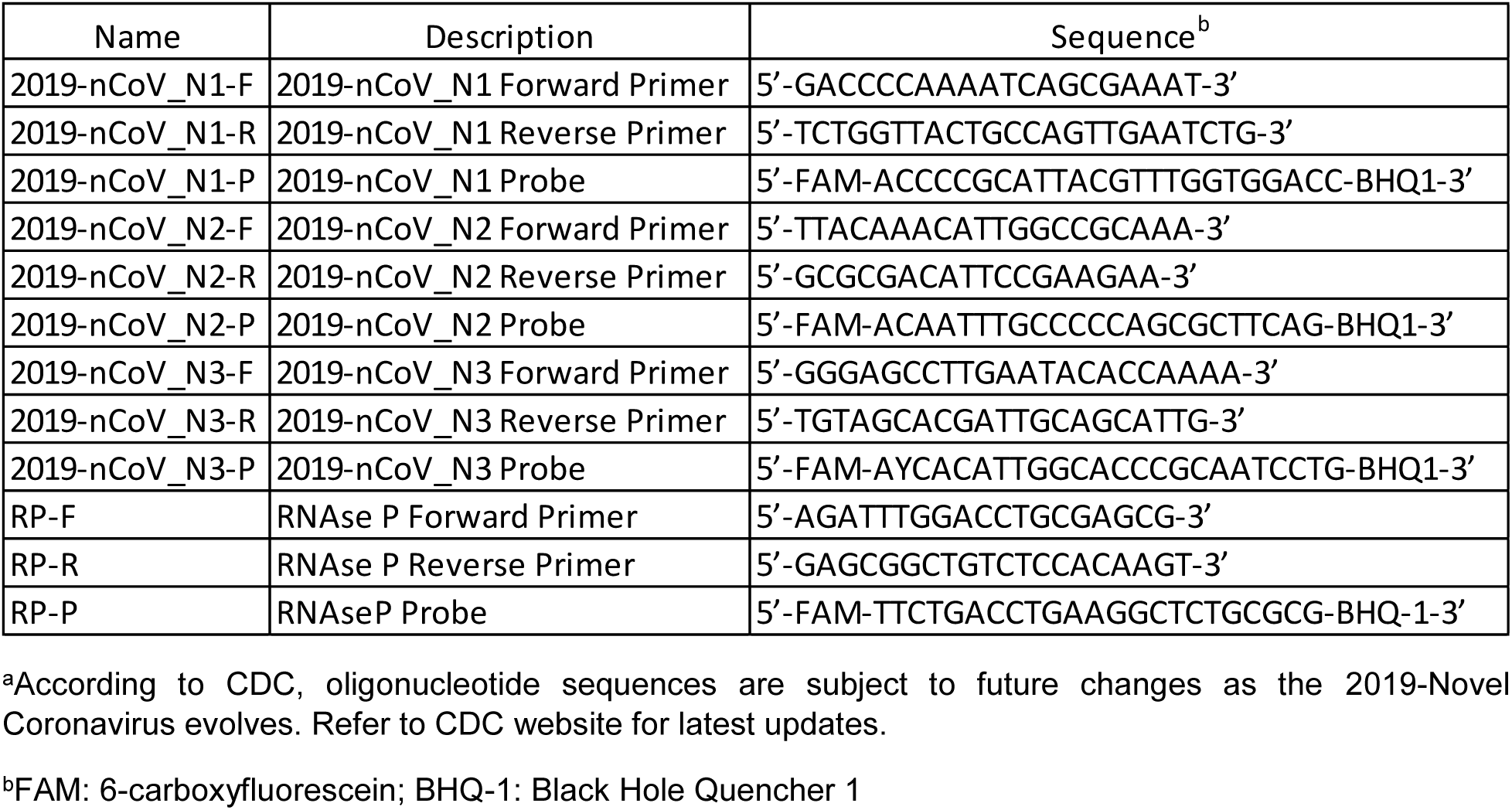
CDC TaqMan RT-qPCR primers and probes for SARS-CoV-2^a^ (adapted from https://www.cdc.gov/coronavirus/2019-ncov/lab/rt-pcr-panel-primer-probes.html).

### Reverse transcription (RT) qPCR assay

RT-qPCR assays were assembled in a total volume of 25 µL containing the indicated buffer at 1X strength (**Table 2**). The buffer was supplemented with 0.4 mM deoxyribonucleotides (dNTP), 402 nM each of forward and reverse PCR primer pairs, 102 nM of the TaqMan probe, and 2.5 units of Taq DNA polymerase from indicated commercial vendors. Indicated copies of SARS-CoV-2 viral genomic RNA, SARS-CoV-2 N gene armored RNA or RNaseP armored RNA prepared in TE buffer (10 mM Tris-HCl, pH 7.5, 0.1 mM EDTA, pH 8.0) immediately prior to use were added to RT-qPCR reactions containing corresponding PCR primers. Negative control reactions did not receive any specific templates. Amplicon accumulation was measured in real-time by incubating the reactions in a LightCycler96 qPCR machine (Roche, Basel, Switzerland) programmed to hold 60 °C for 30 min followed by 95 °C for 10 min prior to undergoing 55 cycles of 95 °C for 15 sec and 60 °C for 30 sec. TaqMan probe fluorescence was measured during the amplification step (60 °C for 30 sec) of each cycle using the FAM channel.

**Table 2.**
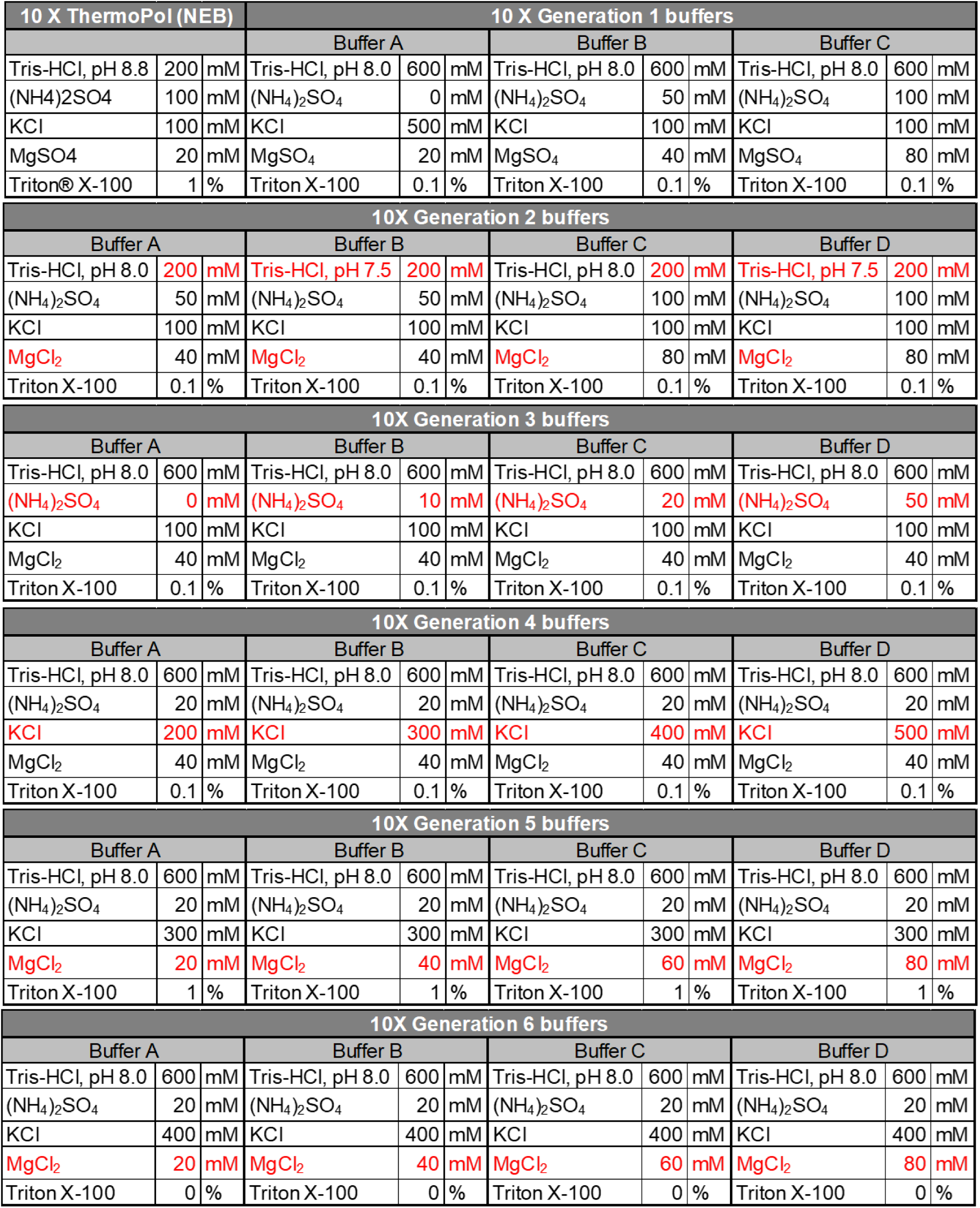
Composition of buffers used in this study.

In some experiments, the initial reverse transcription step (60 °C for 30 min) was eliminated and the reactions were directly subjected to denaturation at 95 °C for 10 min followed by 55 cycles of PCR amplification as described above. In some experiments, RT-qPCR tests were subjected to a heat kill step prior to reverse transcription by incubating the reactions at 95 °C for 5 min. In some experiments, the RNA templates were treated with DNase I prior to RT-qPCR analysis. Briefly, 1 × 10^7^ copies/µL (5×10^8^ total copies) of armored N gene RNA or 2 × 10^3^ copies/µL (10^5^ total copies) of SARS-CoV-2 viral genomic RNA were incubated with 0.5 units of DNase I for 10 min at 37 °C. DNase I was then inactivated by adding EDTA to a final concentration of 5 mM and heating at 65 °C for 10 min.

## RESULTS

### Buffer optimization for Taq DNA polymerase-mediated reverse transcription-qPCR

While Taq DNA polymerase has previously been shown to possess reverse transcriptase activity,^*5, 6*^ it is not commonly thought of as an enzyme that would be readily used for reverse transcription in many assays. To verify this activity and to determine whether Taq polymerase-mediated reverse transcription might be leveraged for single enzyme RNA detection, we carried out the CDC-approved SARS-CoV-2-specific N1 TaqMan RT-qPCR assay using only Taq DNA polymerase and its accompanying commercial reaction buffer, ThermoPol, (New England Biolabs) seeded with different copies of N gene armored RNA (Asuragen), a commercial template preparation that is devoid of DNA. Even though the only polymerase present in these reactions was Taq DNA polymerase (NEB), and no dedicated reverse transcriptase was added, amplification curves were generated in response to 3 × 10^5^, 3 × 10^4^, and 3 × 10^3^ copies of the SARS-CoV-2 N gene armored RNA templates (**Figure 1**).

**Figure 1.**
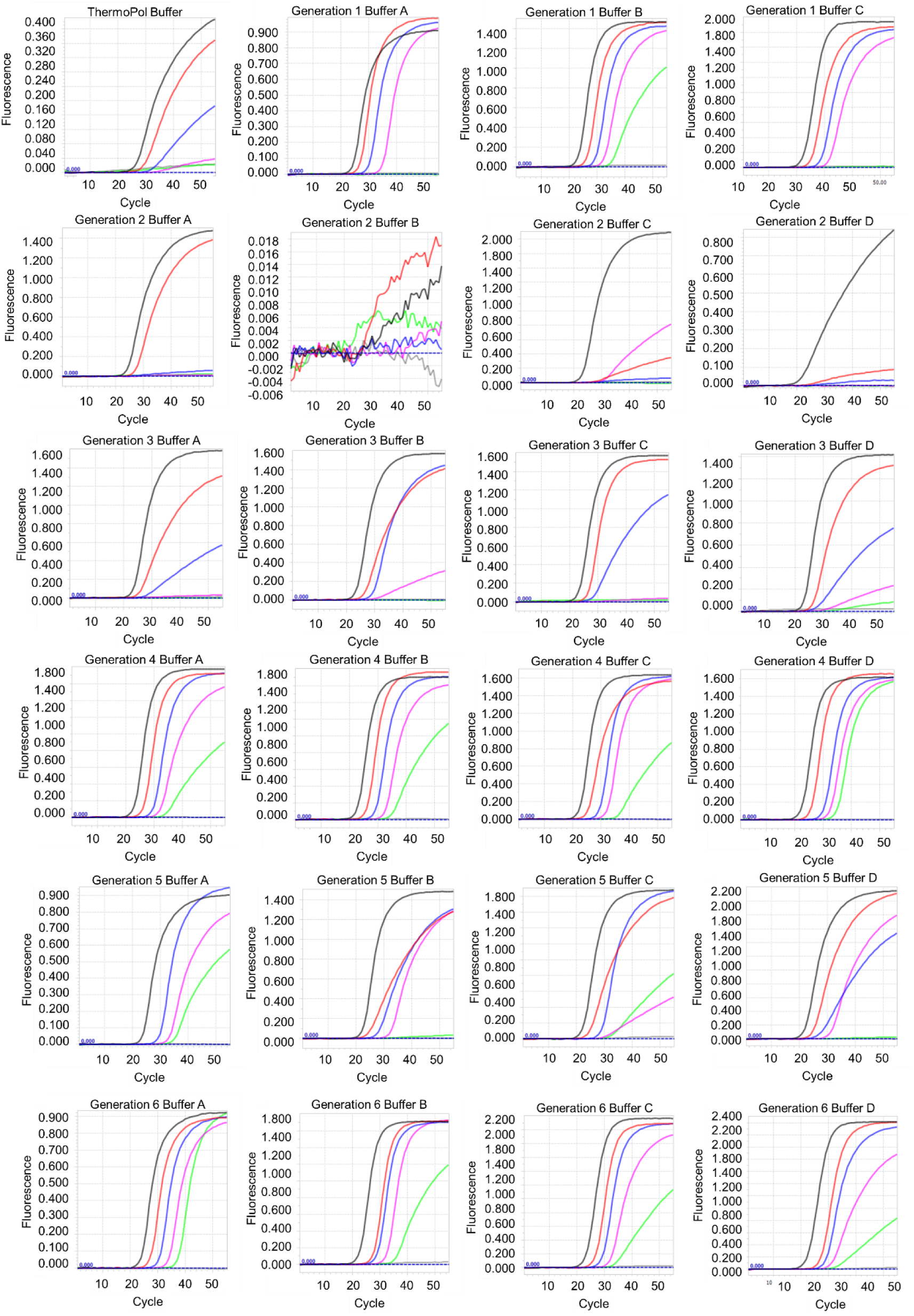
SARS-CoV-2 N1 TaqMan RT-qPCR assays performed using NEB Taq DNA polymerase and N gene armored RNA in indicated buffers. Buffer compositions are detailed in Table 2. Amplification curves resulting from 3 × 10^5^ (black traces), 3 × 10^4^ (red traces), 3 × 10^3^ (blue traces), 3 × 10^2^ (pink traces), 30 (green traces), and 0 (gray) copies of SARS-CoV-2 N gene armored RNA are depicted.

We hypothesized that the buffers in which Taq DNA polymerase is commonly used have been optimized for DNA amplification and likely would not support robust reverse transcription. In addition, previous work had explored optimization of buffer conditions for Taq Stoffel DNA polymerase (i.e. “klenTaq”).^*7*^ We therefore undertook a series of buffer optimizations in which we sequentially varied: buffer pH, Tris concentration, the concentration of monovalent cations ((NH_4_)_2_SO_4_ and KCl) and divalent cations (MgSO_4_), and the concentration of the non-ionic detergent Triton X-100 (**Table 2**). These variables were all chosen based on knowledge of the reaction. The optimum pH for Taq DNA polymerase activity is reported to be between 7.0 and 8.0, with highest activity observed in Tris-HCl-based buffer at pH 7.8,^*8*^ and increasing pH is one of the factors that decreases Taq fidelity.^*9*^ Monovalent cations, such as K^+^, are known to stimulate the catalytic activity of Taq DNA polymerase with optimal activity being observed at about 60 mM KCl,^*8*^ and higher ionic strength can promote primer annealing. Ammonium ions (NH_4_^+^), on the other hand, may have a destabilizing effect, especially on weak hydrogen bonds between mismatched primers and templates. The divalent cation Mg^2+^ is essential for the catalytic activity of Taq DNA polymerase, and its concentration is frequently varied to obtain optimum amplification^*8, 10*^ Finally, the non-ionic detergent Triton X-100 is thought to reduce nucleic acid secondary structure and may influence RT-PCR specificity.

When SARS-CoV-2 N1 TaqMan RT-qPCR assays were performed in the Generation 1 buffers A, B, and C, Taq polymerase generated distinct amplification curves with 10-fold improvement in sensitivity relative to the commercial ThermoPol buffer (**Figure 1**). In fact, in Generation 1, buffer B, amplification curves were observed with as few as 30 copies of armored RNA templates. To identify and hone parameters, we tested the same TaqMan RT-qPCR assay in four Generation 2 buffers with lowered Tris concentrations that matched that of the ThermoPol buffer. In addition, MgSO_4_ was replaced with MgCl_2_, a more commonly used source of Mg^2+^ ions in PCR. However, reduction in Tris concentrations caused a significant drop in RT-qPCR sensitivity compared to both Generation 1 and ThermoPol buffers (**Figure 1**). Therefore, the Tris concentration and pH in 1X buffers were held constant at 60 mM and pH 8.0 from Generation 3 onwards. In Generations 3-6 we sequentially varied the concentration of (NH_4_)_2_SO_4_ from 0 to 5 mM, KCl from 20 to 50 mM, MgCl_2_ from 2 to 8 mM, and Triton X-100 from 0 to 0.1%, and arrived at an optimized Generation 6 buffer A (Gen 6 A) that contained 60 mM Tris, pH 8.0, 2 mM (NH_4_)_2_SO_4_, 40 mM KCl, 2 mM MgCl_2_, and no Triton X-100. Robust amplification curves were generated and as few as 10 copies/µL (30 copies total) of the RNA template could be detected when Gen 6 A buffer was used.

To confirm this novel application of Taq DNA polymerase, we tested two different commercial sources of Taq DNA polymerase, NEB and Thermo Fisher, using additional RNA templates and TaqMan RT-qPCR assays: (i) CDC N1, N2, and N3 TaqMan RT-qPCR assays of SARS-CoV-2 genomic RNA purified from infected cells (ATCC) and (ii) CDC RNaseP TaqMan RT-qPCR assay of RNaseP armored RNA. Similar to our results with armored N gene RNA, the NEB Taq DNA polymerase was able to perform TaqMan RT-qPCR analysis of SARS-CoV-2 viral genomic RNA with all three CDC assays (**Figure 2**). As expected, Gen 6 A buffer improved RT-qPCR performance of Taq DNA polymerase allowing detection of as few as 6 copies of viral RNA in all three N gene assays. The RT-qPCR ability was not restricted to the NEB Taq DNA polymerase. Taq DNA polymerase from a different commercial vendor, Thermo Fisher, could also support RT-qPCR analysis of viral genomic RNA. Similar to the NEB enzyme, Thermo Fisher Taq DNA polymerase also demonstrated better activity in Gen 6 A buffer and was able to detect viral genomic RNA using all three N gene assays, albeit with a higher detection limit (**Figure 2**). Both NEB and Thermo Fisher Taq DNA polymerases were also able to perform TaqMan RT-qPCR analysis of RNaseP armored RNA using the CDC assay (**Figure 2**). These results suggest that Taq DNA polymerase can support TaqMan RT-qPCR analyses of RNA in one-enzyme reactions.

**Figure 2.**
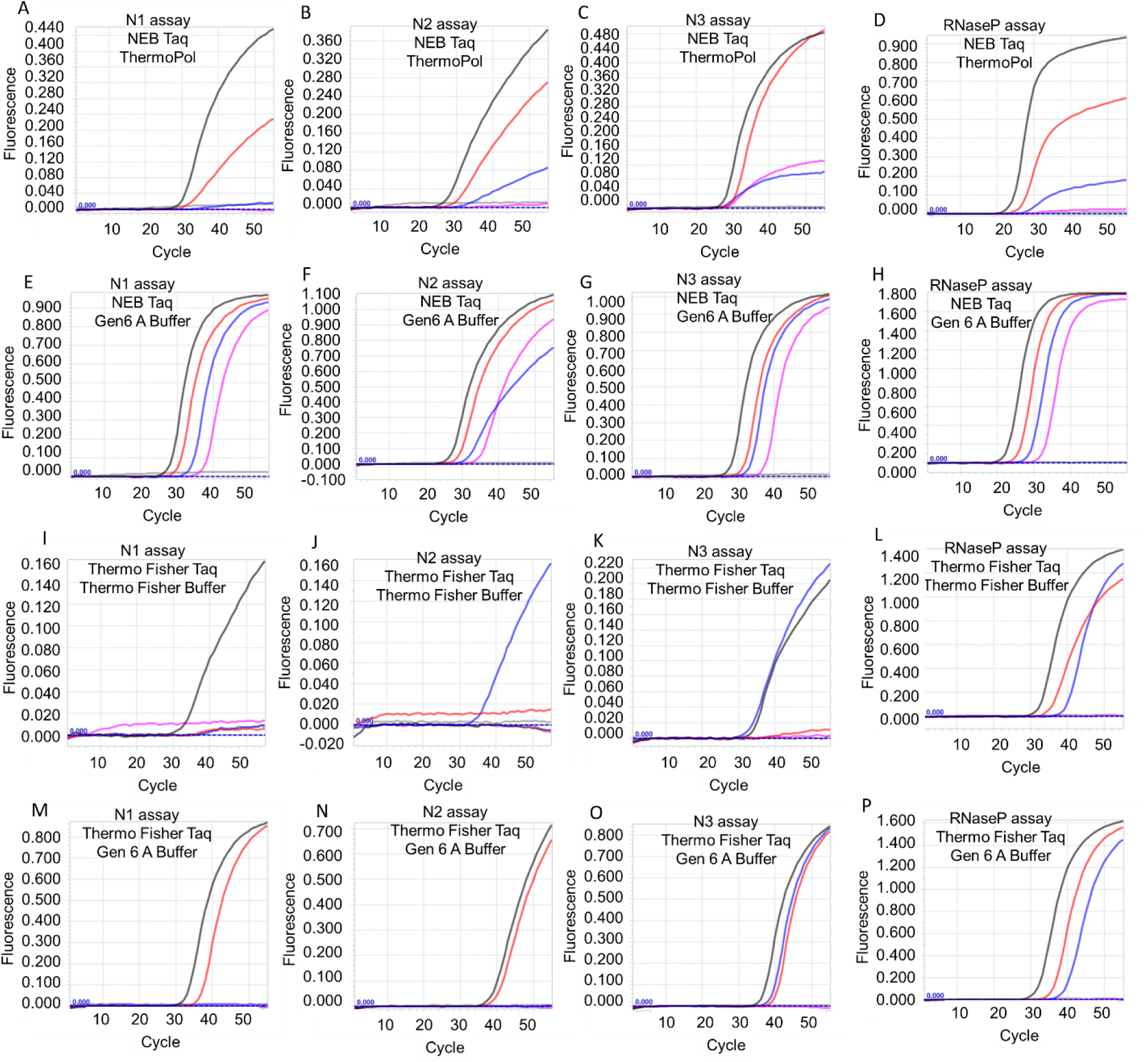
TaqMan RT-qPCR analysis of SARS-CoV-2 viral genomic RNA and RNaseP armored RNA using Taq DNA polymerase-based one-enzyme assays. CDC SARS-CoV-2 N gene assays, N1, N2, and N3, and RNaseP assay were performed using Taq DNA polymerase from either NEB (panels A-H) or Thermo Fisher (panels I-P). Assays were performed either using the companion commercial buffer (panels A-D and panels I-L) or using Gen 6 A buffer (panels E-H and panels M-P). Amplification curves from 6000 (black traces), 600 (red traces), 60 (blue traces), 6 (pink traces), and 0 (gray traces) copies of viral genomic RNA are depicted in panels A-C, E-G, I-K, and M-O. Amplification curves from 3 × 10^5^ (black traces), 3 × 10^4^ (red traces), 3 × 10^3^ (blue traces), 3 × 10^2^ (pink traces) and 0 (gray traces) copies of armored RNaseP RNA are depicted in panes D, H, L, and P.

To further prove that the reverse transcriptase is inherent in Taq polymerase itself, we incubated RT-qPCR assays at 95 °C for 5 min prior to reverse transcription, which should inactivate any contaminating mesophilic reverse transcriptases (**Supplementary Figure 1**). Taq polymerase was still fully capable of RT-qPCR.

### Reverse transcription is necessary for efficient Taq DNA polymerase-mediated RT-qPCR

The armored RNA templates and viral genomic RNA templates used in these studies are theoretically devoid of DNA templates. To demonstrate that the TaqMan RT-qPCR amplification signals generated by Taq DNA polymerase are not due to amplification of contaminating DNA templates, we treated the RNA templates with DNase I prior to RT-qPCR amplification. As shown in **Figure 3**, Taq DNA polymerase-mediated TaqMan RT-qPCR assays generated amplification signals from both DNase treated genomic RNA and armored RNA. This was true of not only NEB and Thermo Fisher Taq DNA polymerase but also a preparation of Taq DNA polymerase purchased from Promega (data not shown). While N2 and N3 assays could still detect the smallest template quantities tested - 6 copies of viral genomic RNA or 30 copies of armored N RNA – most amplification curves suffered a small increase of about 1-2 Ct in their time to detection (**Figure 3**). In contrast, the N1 assay demonstrated anywhere between a 1 and 5 Ct delay with DNase treatment of RNA templates, and failed to generate signal from the lowest template concentrations.

**Figure 3.**
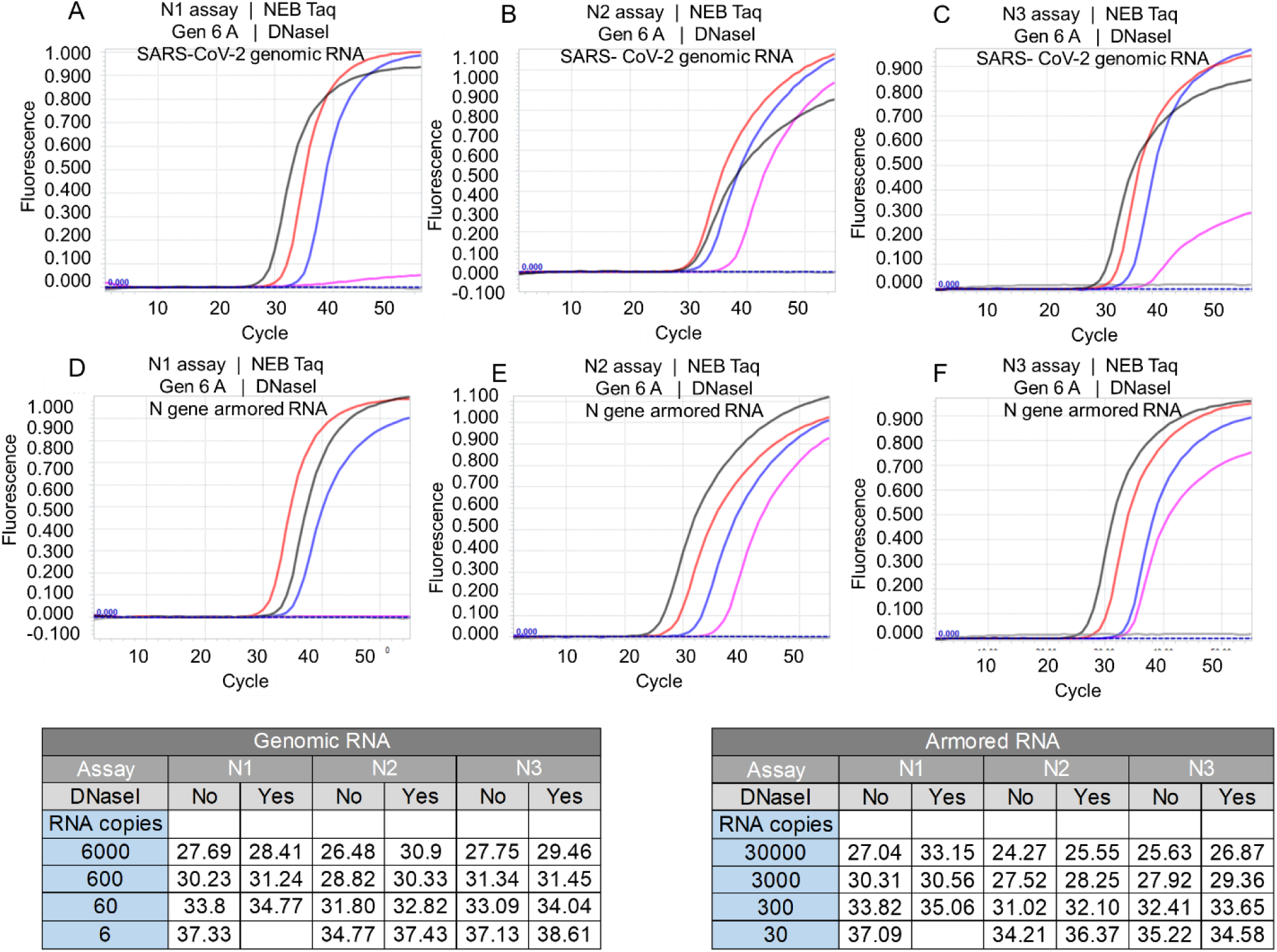
Effect of DNase I treatment on Taq DNA polymerase-mediated RT-qPCR assay. Taq DNA polymerase purchased from NEB was used to operate CDC SARS-CoV-2 N1, N2, and N3 TaqMan RT-qPCR assays using SARS-CoV-2 viral genomic RNA (panels A-C) or N gene armored RNA (panels D-F) treated with DNase I. Amplification curves shown in panels A-C resulted from 6000 (black traces), 600 (red traces), 60 (blue traces), 6 (pink traces), and 0 (gray traces) copies of SARS-CoV-2 genomic RNA. Amplification curves in panels D-F resulted from 30,000 (black traces), 3,000 (red traces), 300 (blue traces), 30 (pink traces) and 0 (gray traces) copies of N gene armored RNA. Representative Ct values for RT-qPCR amplification of indicated copies of untreated and DNase I treated SARS-CoV-2 genomic RNA and N gene armored RNA are tabulated.

To confirm that reverse transcription step in the TaqMan RT-qPCR assay was indeed necessary in order to generate amplification curves in response to template RNA, we executed CDC SARS-CoV-2 N1, N2, and N3 assays using NEB or Thermo Fisher Taq DNA polymerases without performing the reverse transcription step prior to qPCR amplification. Eliminating a 30 min incubation at 60 °C (the putative reverse transcription step) prior to PCR thermal cycling also eliminated robust amplification of viral RNA templates (**Supplementary Figure 2**). The TaqMan probe signal remained at background levels in all reactions except the N3 assays performed in Gen 6 A buffer. Even in this case, amplification curves were significantly delayed and only apparent in the presence of relatively high numbers of RNA templates. For instance, Taq obtained from NEB detected 6000 and 600 copies of viral genomic RNA (with Ct values of 43.42 and 46.55, respectively). These same copy numbers of template RNA when subjected to a Taq-mediated reverse transcription step prior to qPCR amplification typically yielded Ct values of 27 and 30 (**Figure 2**). These results demonstrate that Taq polymerase can generate amplicons from RNA during a normal thermal cycling reaction, and that a pre-incubation step greatly improves detection.

## CONCLUSIONS

These results suggest that, given the correct buffer conditions, Taq DNA polymerase can perform reverse transcription and TaqMan qPCR in one-pot reactions. While the use of a proficient reverse transcriptase along with a proficient thermostable DNA polymerase may still be the best option for many applications, the ability to use only a single enzyme in assays for the detection of SARS-CoV-2 RNA opens the way to new diagnostic approaches, especially in resource-poor settings where reagent availability or production may be issues. However, it is also evident from our experiments that the variability in amplification efficiency observed as buffer conditions were iteratively improved may mean that similar buffer optimizations will need to be routinely carried out with other templates. It is nonetheless possible that the reverse transcriptase activity of Taq DNA polymerase, once fully optimized, will prove useful in many different applications, up to and including those where many different templates are present, such as RNASeq.

## ACKNOWLEDGEMENTS

This work was supported by grants from the Welch Foundation (F-1654), the National Institutes of Health (1-R01-EB027202-01A1), and the National Aeronautics and Space Administration (NNX15AF46G).

