## Supplementary Figures for "One enzyme reverse transcription qPCR using Taq DNA polymerase"

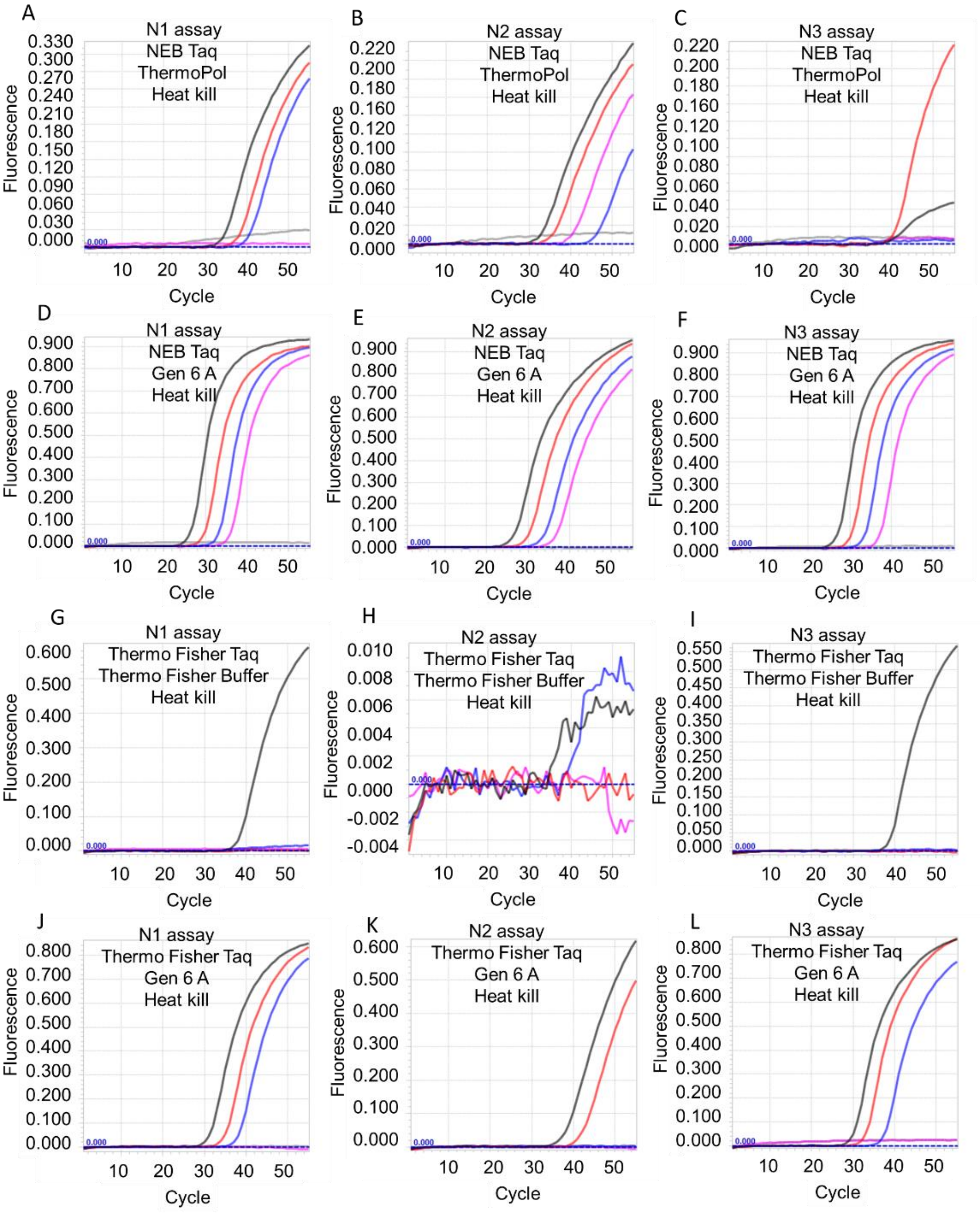


**Supplementary Figure 1.** Effect of heat kill step prior to reverse transcription on Taq DNA polymerase TaqMan RT-qPCR assays. CDC SARS-CoV-2 N1, N2, and N3 TaqMan RT-qPCR assays were setup using NEB (panels A-F) or Thermo Fisher (panels G-L) Taq DNA polymerase and their companion commercial buffers or Gen 6 A buffer. Amplification curves resulting from 6000 (black traces), 600 (red traces), 60 (blue traces, 6 (pink traces), and 0 (gray traces) copies of SARS-CoV-2 genomic RNA are depicted.

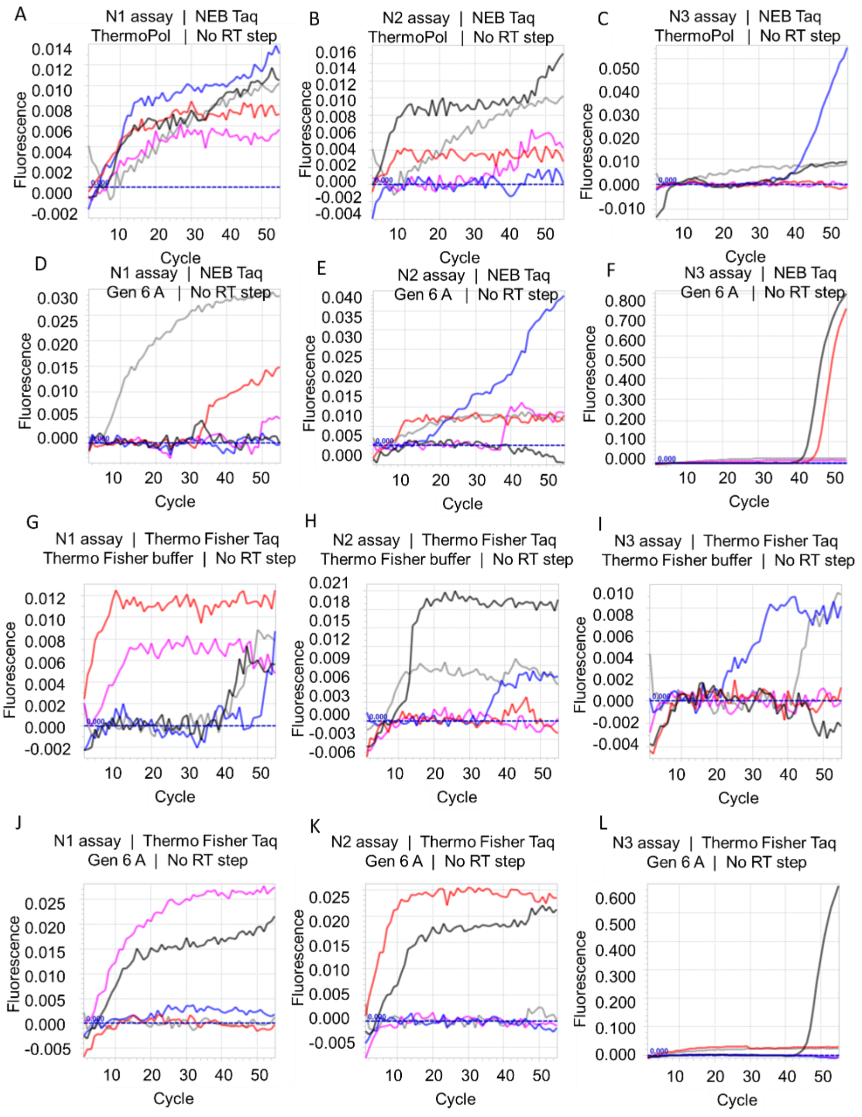


**Supplementary Figure 2.** Effect of eliminating the reverse transcription step prior to qPCR cycling on Taq DNA polymerase-mediated TaqMan RT-qPCR assays. CDC SARS-CoV-2 N1, N2, and N3 TaqMan RT-qPCR assays were setup using NEB (panels A-F) or Thermo Fisher (panels G-L) Taq DNA polymerase and their companion commercial buffers or Gen 6 A buffer. Amplification curves resulting from 6000 (black traces), 600 (red traces), 60 (blue traces, 6 (pink traces), and 0 (gray traces) copies of SARS-CoV-2 genomic RNA are depicted.
